# *Escherichia coli* acid phosphatase AphA expression is upregulated under carbon and phosphate starvations and inhibited by CytR

**DOI:** 10.1101/2023.12.05.570089

**Authors:** Kristina Kronborg, Yong Everett Zhang

## Abstract

This study investigates the regulation of *E. coli aphA* expression under nutrient starvation. Using transcriptional reporters with truncated *aphA* promoter sequences, we found that starvation of carbon and phosphate, but not amino acid, stimulated *aphA* expression through distinct promoter regions. Deletions of *crp* or *cyaA* reduced *aphA* expression, confirming their importance in *aphA* regulation during carbon starvation. Conversely, CytR deletion increased *aphA* expression, suggesting CytR’s role as a repressor of *aphA* expression. Collectively, these data imply a potential connection between CytR, *aphA* expression, and the broader context of natural competence evolution and bacterial nutrient absorption.

## Introduction

Cyclic AMP (cAMP) is a universal signaling molecule important to both eukaryotes and prokaryotes (1,2) and proteins binding to cAMP are thus of high interest. Recently, we screened the *Escherichia coli* proteome and identified the non-specific acid phosphatase AphA as the other target protein of cAMP besides CPR (3). AphA is localized in the periplasmic space where it serves as a scavenging enzyme dephosphorylating a wide range of organic phosphomonoesters that otherwise cannot cross the inner membrane (4,5). Thereby, AphA allows *E. coli* to utilize these various phosphomonoesters for nutritional purposes and hence aid in bacterial survival. Similarly, AphA homolog in *Salmonella typhimurium* facilitates its uptake of external NAD (6). We recently found that in *H. influenzae*, AphA, together with NadN and eP4, serves to coordinate nutritional growth with the bacterium’s competence development (3). *E. coli* is, however, not naturally competent and AphA may play a different role in this bacterium. Previously, it is shown that the expression of *aphA* is induced by cAMP-CRP (7). In this study we aimed to investigate the expression of AphA in *E. coli*, to identify the promotors responsible for *aphA* expression during different starvation conditions, to confirm the requirement of cAMP-CRP for *aphA* expression, and finally to identify regulators of its expression.

## Results

### Carbon and phosphate starvations induce transcription of *E. coli aphA*_*Ec*_

The expression of *E. coli aphA*_*Ec*_ is stimulated by carbon starvation (7). To confirm this observation and to study how *aphA*_*Ec*_ transcription is regulated, we constructed four transcriptional reporter plasmids by cloning the serially truncated promoter regions (P1 to P4 from -433 to +30, **Figure** 1A-C) of *aphA*_*Ec*_ upstream of the *lacZYA* operon of the low-copy-number vector pGH254 previously used to analyze gene expression in *E. coli* (8). We first grew the *E. coli* strains with these reporters in MOPS synthetic rich medium and then shifted the cells to MOPS medium lacking either carbon, amino acid, or phosphate, to quantitate the transcription level of *aphA*_*Ec*_. Consistent with previous report (7), carbon starvation led to an abrupt increase of *aphA*_*Ec*_ transcription from the promoter region P1 and P2, but moderately from P3, and not from P4 (**Figure** 1D), suggesting the -200 to -123 region is critical for *aphA*_*Ec*_ transcription during carbon starvation. A downshift of the reporter strains to low phosphate (from 1.32 mM to 0.067 mM) MOPS medium elicited a slow and moderate increase of *aphA*_*Ec*_ transcription from P2 and P3, but not from P4 or P1 (**Figure** 1E), suggesting that phosphate downshift stimulates the *aphA*_*Ec*_ transcription as well. At last, a downshift to MOPS medium without amino acid revealed no change of *aphA*_*Ec*_ transcription (**Figure** 1F), suggesting that amino acid starvation, and thus the stringent response, has no effect in *aphA*_*Ec*_ transcription. Of note, P2 and P3, but not P1, are generally important for high basal level of *aphA* expression before nutrient downshift (**Figure** 1D-1F). Altogether, these data suggest that carbon and phosphate starvations induce the expression of *aphA*_*Ec*_ via overlapping but distinct promoter regions. Cleavage of nucleotides by AphA releases nucleoside and phosphate, serving as carbon and phosphate sources respectively. The data is consistent with the notion that AphA_Ec_ may be involved in bacterial utilization of nucleotides as both carbon and phosphate sources. The coupled stimulation of *aphA*_*Ec*_ transcription upon either carbon or phosphate downshift thus connects AphA_Ec_ to the nucleotide utilization in *E. coli*.

**Figure 1.**
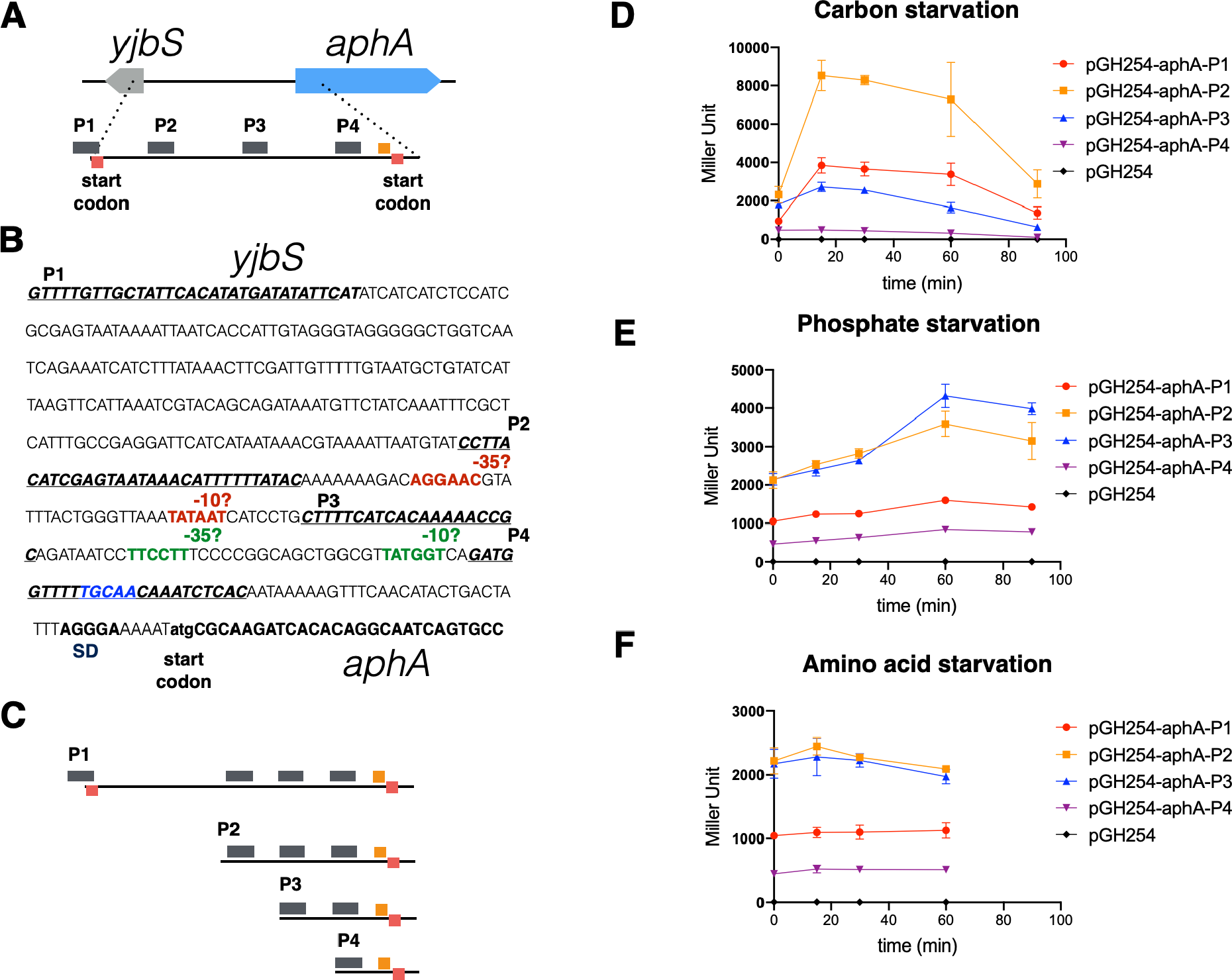
Carbon and phosphate starvation stimulate the expression of *E. coli aphA*. (**A**) Schematic overview showing the promoter region of *E. coli aphA* gene, with the start codons and potential four promoter sequences marked (bold, italic). (**B**) Sequence annotation of the promoter region of *E. coli aphA*. Text highlighted in blue is the CytR box. (**C**) Schematic overview showing the promoter regions of *aphA*, which were transcriptionally fused to a *lacZ* reporter gene in the plasmid pGH254 (8). (**D, E, F**) Beta-galactosidase assays of the various transcriptional reporters during starvations of carbon, phosphate, and amino acid. Two biological replicates were performed, and the average and standard errors are shown.

### CyaA and CRP are essential for the *aphA*_*Ec*_ expression

Carbon starvation in *E. coli* activates the cAMP-dependent CRP regulon. To confirm the link between carbon starvation and *aphA*_*Ec*_ transcription, we transformed the reporter plasmid pGH254-*aphA*-P2 into the KEIO deletion strains (9) of CRP or CyaA and performed a beta-galactosidase assay (see method). Besides, we also included several KEIO strains deleted with other well studied transcriptional regulators, i.e. Sxy, CytR, RpoS, PhoB, and PurR, given their (plausible) connections to bacterial nucleotide metabolism and AphA production in different organisms (10). Transcription of *aphA*_*Ec*_ dropped to basal levels in both Δ*cyaA* and Δ*crp* strains (**Figure** 2), demonstrating their essential roles in *aphA*_*Ec*_ expression upon carbon starvation. Additionally, in the Δ*cytR* mutant strain, transcription of *aphA* increased versus the wild type strain, suggesting that CytR suppresses the *aphA*_*Ec*_ transcription (see Discussion below). Indeed, inspection of the promoter region revealed a potential CytR box (TGCAA, TTGC/tA) (11) within the P4 and Nt of the *aphA*_*Ec*_ gene. The other mutant strains tested showed marginal, if at all, decrease of the *aphA*_*Ec*_ transcription. These data indicate that the production of AphA_Ec_ is controlled by both carbon and nucleotides metabolism.

**Figure 2.**
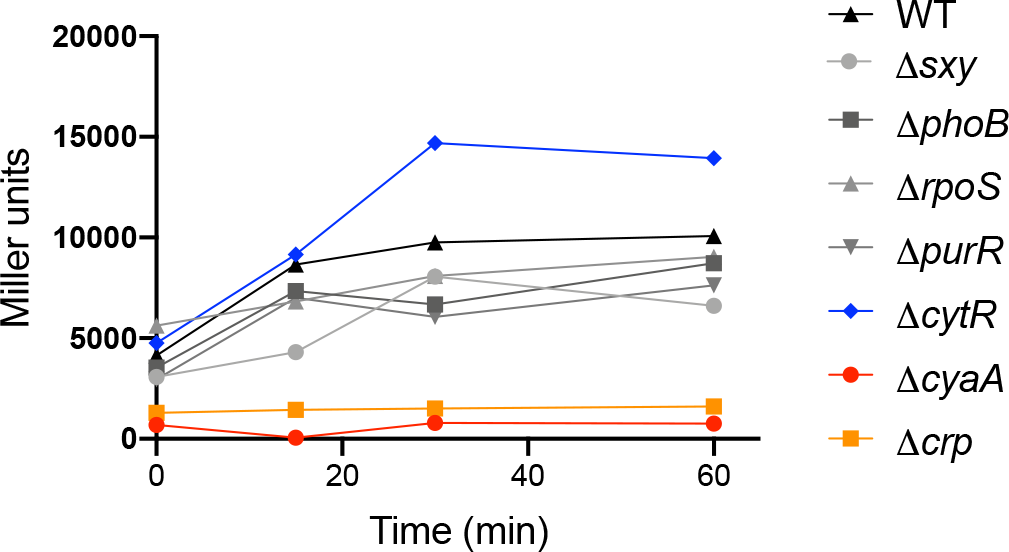
Beta-galactosidase assays during carbon starvation of various mutant strains of *E. coli*. Time zero indicates the time right before the downshift of cells from the MOPS complete medium to the same medium depleted of glucose. Two biological replicates were performed, and one representative curve is shown.

## Discussion

This short study of *aphA* expression in *E. coli* confirms the requirement for cAMP-CRP for *aphA* expression. It furthermore suggests CytR to be a repressor of *aphA* expression. Instead, other tested regulators appear to exert only a minimal impact on *aphA* expression under carbon starvation. CytR typically represses gene expression by interacting with CRP and blocking the binding site of CRP (12). Upon binding to cytidine, the inducer, CytR unmasks the binding site of CRP, leading to derepressed gene expression. The discovery of a potential CytR box in the *aphA* promoter region (**Figure** 1B) is consistent with this notion and the observed increased expression of *aphA* (**Figure** 2). Intriguingly, CytR is required for competence development in *Vibrio cholerae* (13). Recently, the introduction of external cytidine (100 mM) diminished *V. cholerae* competence to a level comparable to the deletion of *cytR*_Vc_ (14). On the other hand, purine nucleotides are involved in the competence development of *Haemophilus influenzae* (15) (10). Altogether, these data suggest a potential link among CytR, *aphA* expression, and the broader context of natural competence evolution and bacterial nutrient absorption.

### Experimental procedures

#### Construction of transcriptional reporters of aphA-lacZ

To make the APHA-1 transcriptional reporter, the primers pYZ262 and pYZ266 were used to amplify the P1 promoter region (Figure 1) of *aphA*_*Ec*_ and cloned into the plasmid vector pGH254 (8) via the EcoRI and BamHI restriction sites. This places the P1 promoter region of *aphA*_*Ec*_ in front of the *lacZYA* genes on pGH254 resulting in the transcriptional reporter pGH254-APHA-1. Likewise, P2, P3 and P4 promoter regions of *aphA*_*Ec*_ were amplified using primers pYZ263 and pYZ266, primers pYZ264 and pYZ266, and primers pYZ265 and pYZ266, respectively, and cloned into pGH254 to construct the pGH254-APHA-2, pGH254-APHA-3, pGH254-APHA-4, respectively.

#### Beta-galactosidase assay

Overnight cultures of *E. coli* strains grown in MOPS medium (18 h, 37°C, 160 rpm) were back-diluted into fresh MOPS medium and grown to OD_600_=0.3-0.5 before they were shifted to a MOPS medium lacking either phosphate, glucose, or amino acids. During growth, pelleted cell samples were collected and immediately frozen. Next, samples were thawed and resuspended in a resuspension buffer composed of 100 mM Tris (pH 7.8), 32 mM Na_3_PO_4_ (pH 7.8), 1 mM PMSF and 19 mM β-mercaptoethanol and were sonicated (10% amplitude) until complete lysis was observed. Protein concentration was measured using the Bradford assay, and equal amounts of total protein from each lysed sample were resuspended in a reaction buffer composed of 76 mM Tris (pH 7.8), 24 mM Na_3_PO_4_ (pH 7.8), 0.03% SDS, 1.3 mM MgSO_4_, 0.4 mg/ml DNase I, 1 mM PMSF and 19 mM β-mercaptoethanol to a final volume of 75 μl. The β-galactosidase reaction was started by adding 60 μl 4 mg/ml ONPG to the reaction mixture and was terminated by the subsequent addition of 150 μl 1 M Na_2_CO_3_. Finally, the samples were spun down for 2 min at 14600 rpm and 200 μl supernatant was transferred to a clear flat-bottom 96-well plate (Greiner) and the absorbance was read at 420 nm in a plate reader.

**Table 1:** Bacterial strains used in this study.

| Strain | Relevant features | Reference |
| --- | --- | --- |
| YZ351 | TB28 pGH254-APHA-1, Kan | This study |
| YZ352 | TB28 pGH254-APHA-2, Kan | This study |
| YZ353 | TB28 pGH254-APHA-3, Kan | This study |
| YZ354 | TB28 pGH254-APHA-4, Kan | This study |
| YZ356 | TB28 pGH254, Kan | Laboratory stock |
| YZ1047 | Keio *sxy::kan*, pGH254-APHA-2, Kan, Cam | This study |
| YZ1048 | Keio *phoB::kan*, pGH254-APHA-2, Kan, Cam | This study |
| YZ1049 | Keio *rpoS::kan*, pGH254-APHA-2, Kan, Cam | This study |
| YZ1050 | Keio *purR::kan*, pGH254-APHA-2, Kan, Cam | This study |
| YZ1051 | Keio *cytR::kan*, pGH254-APHA-2, Kan, Cam | This study |
| YZ1052 | Keio *cyaA::kan*, pGH254-APHA-2, Kan, Cam | This study |
| YZ1053 | TB28 *crp::tet*, pGH254-APHA-2, Tet, Cam | This study |
| YZ1054 | Keio WT, pGH254-APHA-2, Cam | This study |
| YZ1055 | Keio *sxy::kan*, pGH254-APHA-3, Kan, Cam | This study |
| YZ1056 | Keio *phoB::kan*, pGH254-APHA-3, Kan, Cam | This study |
| YZ1057 | Keio *rpoS::kan*, pGH254-APHA-3, Kan, Cam | This study |
| YZ1058 | Keio *purR::kan*, pGH254-APHA-3, Kan, Cam | This study |
| YZ1059 | Keio *cytR::kan*, pGH254-APHA-3, Kan, Cam | This study |
| YZ1060 | Keio *cyaA::kan*, pGH254-APHA-3, Kan, Cam | This study |
| YZ1061 | TB28 *crp::tet*, pGH254-APHA-3, Tet, Cam | This study |
| YZ1062 | Keio WT, pGH254-APHA-3, Cam | This study |

**Table 2:**
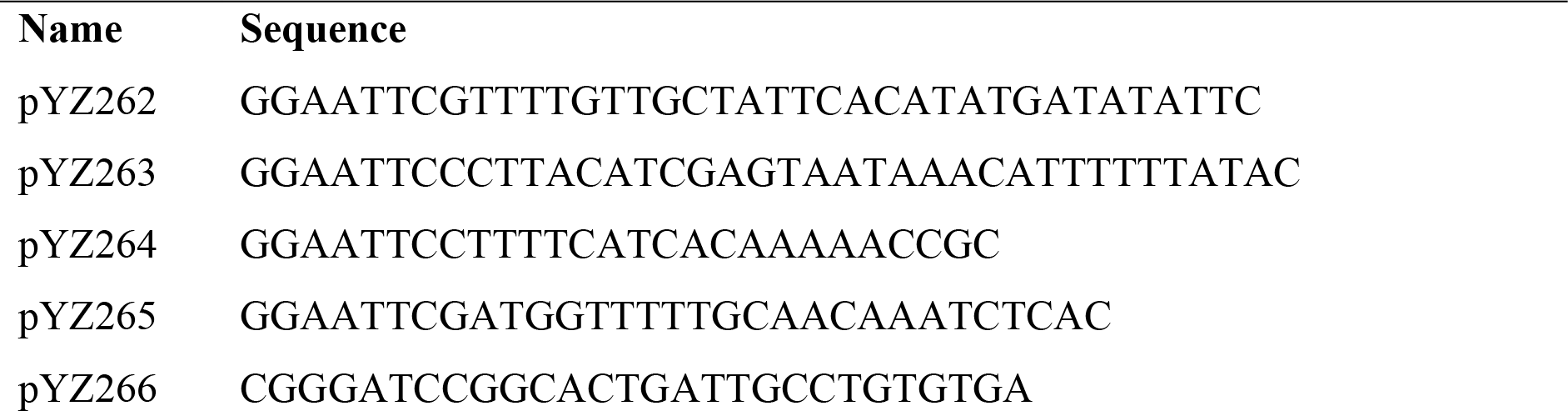
Primers used in this study Name Sequence.

## Acknowledgement

We are grateful to Kenn Gerdes for useful discussions. This study was supported by a Novo Nordisk Foundation Project Grant (NNF19OC0058331) to Y.E.Z.

